# A multivariate network analysis of ring- and diffuse-porous tree xylem vasculature segmented by convolutional neural networks

**DOI:** 10.1101/2023.01.10.523508

**Authors:** Annika Erika Huber, Mohammad Haft-Javaherian, Maxime Berg, Asheesh Lanba, Sylvie Lorthois, Taryn L. Bauerle, Nozomi Nishimura

## Abstract

The xylem network, the water conduction system in wood determines the ability of trees to avoid hydraulic failure during drought stress. The capability to withstand embolisms, disruptions of the water column by gas bubbles that contribute to hydraulic failure, is mainly determined by the anatomical arrangement and connectedness (topology) of xylem vessels. However, the quantification of xylem network characteristics has been difficult, so that relating network properties and topology to hydraulic vulnerability and predicting xylem function remains challenging. We studied the xylem vessel networks of three diffuse- (*Fagus sylvatica, Liriodendron tulipifera, Populus x canadensis*) and three ring-porous (*Carya ovata, Fraxinus pennsylvatica, Quercus montana*) tree species using volumetric images of xylem from laser ablation tomography (LATscan). Using convolutional neural networks for image segmentation, we generated three-dimensional, high-resolution maps of xylem vessels, with detailed measurements of morphology and topology. We studied the network topologies by incorporating multiple network metrics into a multidimensional analysis and simulated the robustness of these networks against the loss of individual vessel elements that mimic the obstruction of water flow from embolisms. This analysis suggested that networks in *Populus x canadensis* and *Carya ovata* are quite similar despite being different wood types. Similar networks had comparable experimental measurements of P50 values (pressure inducing 50% hydraulic conductivity loss) obtained from hydraulic vulnerability curves, a common tool to quantify the cavitation resistance of xylem networks. This work produced novel data on plant xylem vessel networks and introduces new methods for analyzing the biological impact of these network structures.

**Significance statement:** The resilience of fluid transport networks such as xylem vessels that conduct water in trees depends on both the structure of the network and features of the individual network elements. High-resolution reconstruction of xylem networks from six tree species provided novel, three-dimensional, structural data which enabled the xylem networks to be described using graph theory. Using an array of network metrics as multidimensional descriptors, we compared the xylem networks between species and showed relationships to simulated and experimental measures of drought resistance. In addition to providing insight on drought resistance, these approaches offer new ways for comparative analysis of networks applicable to many systems.

## Introduction

Across biological organisms, survival depends on reliable fluid transport in vascular networks such as xylem networks in plants and blood vessels in animals. In such fluid transport networks, topology impacts hydraulic transport efficiency, dictates the distribution of fluid throughput, and determines vulnerability in response to disruptions (1–3). Representing a network in a graph facilitates quantitative descriptors and the analysis of functional efficiency using many different methods (4–9). Therefore, the ability to accurately capture biological networks as graphs is a crucial step toward understanding the robustness of these complex systems. However, despite recent advances in imaging technology that allows for xylem network imaging, technical challenges remain in mapping, analyzing, and thus comparing large networks across tree species (10). As a result, to date, models based on statistical properties or hypothesized structures of networks are the primary tool used to understand how network connectivity affects complex traits such as xylem network vulnerability (6, 7, 11, 12). Here, we developed a novel approach to mapping xylem networks of six different tree species representing two different wood types (diffuse versus ring porous) and explored methods to objectively compare xylem networks.

In trees, water flows along a negative water potential (tension) gradient between the soil and subsaturated air through a network of interconnected xylem vessels (xylem conduits) (3). Xylem vessel diameters range from 5-400 μm and have random (diffuse-porous) or bimodal (adding large diameter conduits in the spring and small diameter in the latewood; ring-porous) distributions within one tree growth ring. Xylem vessels are constructed by end-to-end aligned vessel elements, that at maturity consist of secondary cell walls left after cell death, and connected via perforated or dissolved end walls that provide a low resistance pathway for vertical water flow. Adjacent xylem vessels are horizontally connected by intervessel pit connections that consist of many pit pores permeabilizing the vessel wall (13). Pit pores are many-fold smaller than xylem vessels, ranging from 1-10 nm in diameter, but because of their large numbers can contribute substantially to water conductance (14, 15). Collectively, both xylem vessel topology and intervessel pit connections are important components of the water transport efficiency through xylem networks (16).

Although intervessel pit pores are beneficial for water transport they can also become a liability during drought. During water shortages, the tension in the water column increases and water can spontaneously undergo a rapid phase change from the fluid to the vapor state (cavitation). The embolisms formed during cavitation by the expansion of microscopic air nuclei can then expand to block the water flow through xylem vessels (17). Because intervessel pit connections are not only permeable to water but also to air, embolisms can, under increasing tension, be aspirated through pits into neighboring conduits and spread through the xylem networks (18). Consequently, connected networks, with many pit connections, could make networks more vulnerable (9). However, it is also possible that the redundancy of multiple paths for water flow provided by pit connections facilitates the redirection of the water flow around embolism events, making the network more robust against conductance loss (32). These two contrary arguments highlight the necessity for characterizing the architecture of xylem networks to identify network traits affecting embolism resistance.

A common metric for comparing drought tolerance across plant species is the hydraulic vulnerability curves (percent loss of conductance curves; PLC curves) (19–21). P_50_ values describe the water potential corresponding to a 50% hydraulic conductance loss caused by embolism events in the xylem network and can range from −0.2 to −14.0 MPa in trees, with more negative values corresponding to greater drought tolerance (22–24). P_50_ values are an empirical measurement and multiple attempts have been made to correlate anatomical characteristics of xylem vessels with embolism resistance (25–29). However, xylem vessel characteristics alone do not determine the overall hydraulic efficiency or embolism resistance of xylem networks, because hydraulic efficiency and embolism resistance are collective properties of the whole, threedimensional network.

In this study, we investigated the network characteristics that influence hydraulic efficiency and embolism resistance of xylem vessel networks using laser ablation tomography (LATscan), a method that produces sequential high-resolution two-dimensional image stacks to reconstruct three-dimensional networks (30). We mapped three-dimensional xylem networks of three ring-porous and three diffuse-porous tree species. Comparing these two groups is interesting because ring-porous tree species have a higher maximum hydraulic conductance than diffuse-porous tree species due to the larger vessels in the early growth periods of the season in comparison to diffuse-porous tree species (31, 32). However, these large early-season vessels have been reported to be particularly vulnerable to embolism events (29, 33, 34). We investigated the xylem vessel topology of the two wood types by generating xylem network graphs from deep neural network image binarization previously developed to study brain vascular networks (35–37). We then simulated the robustness of the water flow in the network against blocked vessels and correlated these findings to experimental P_50_ values (22, 28, 38). This new approach to studying the xylem network gives new insight into how xylem vessel topology and connectedness affect network robustness to cavitation under drought stress in both ring- and diffuse-porous tree species and sheds light on the structure to function relationship of both ring- and diffuse-porous xylem networks.

## Results

### Network representations from serial sections of branches

Two-to seven-year-old branches from three individual trees of three diffuse-porous and three ring-porous tree species were harvested (Fig 1A). We segmented xylem vessels from 1000 to 5000 laser ablation tomography (LAT) images from the branches (Fig 1B) and then extracted their network structures as graphs. Graphs are mathematical representations that describe a set of relationships (edges) between a set of objects (vertices) and facilitate quantitative analysis of the objects in the context of their relationships. We used multiple graph representations to study and characterize xylem networks using network analysis and fluid mechanics. Because the LAT resolution was too coarse to visualize intervessel pit membranes, we approximated the location of intervessel connections as regions where the distance of the wall between xylem vessels was reduced below a threshold determined for each species as follows. We measured the maximum distance between xylem vessels with intervessel pit connections in scanning electron microscopy images in each species and set this distance as the threshold for locating intervessel pit connections in LAT images (Table 1 and Fig 1C). The diameters and centerlines (Fig 1D) of xylem were extracted from each image.

**Fig 1.**
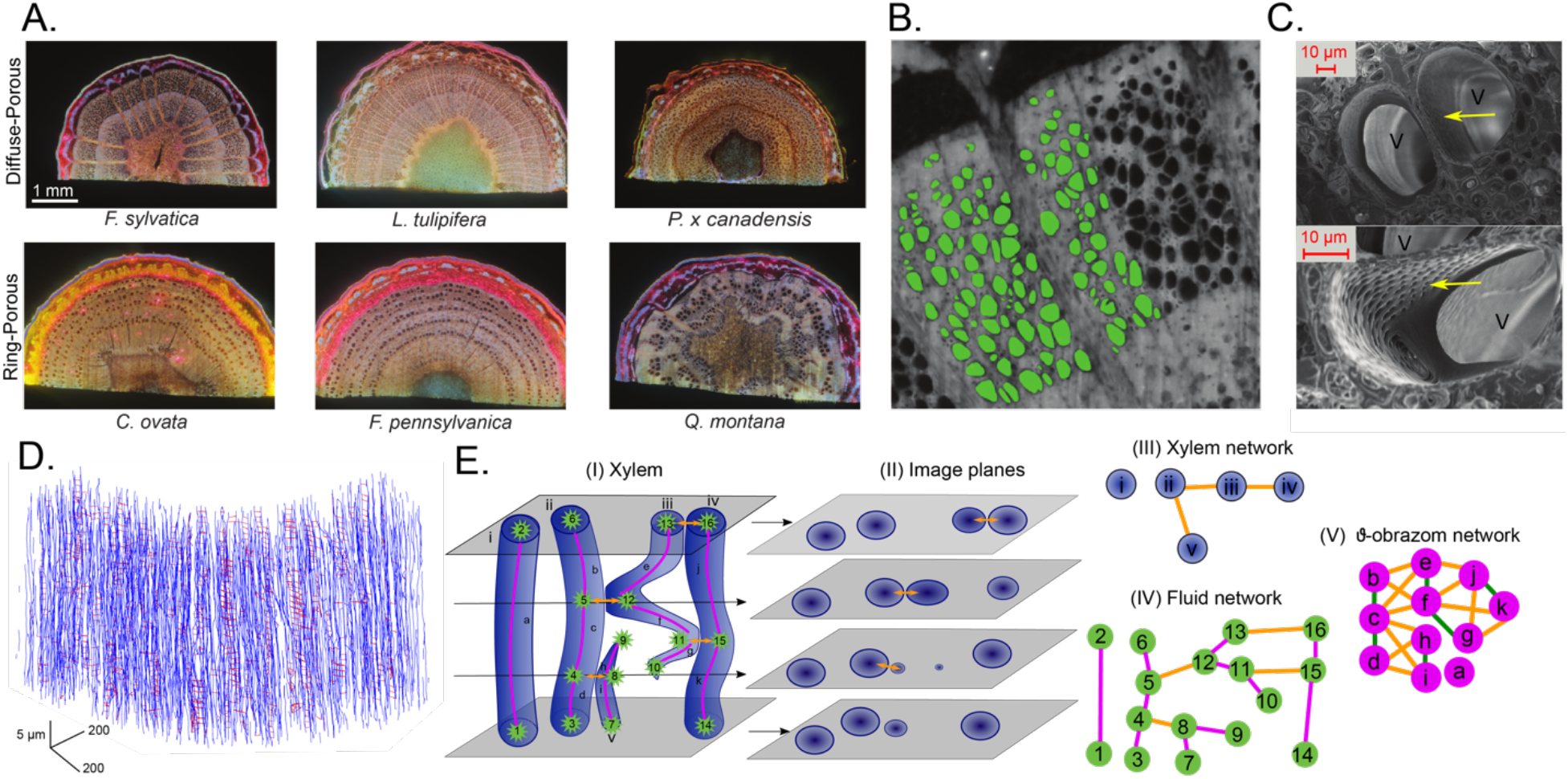
Xylem vessel network representations and the image processing pipeline. A. Example laser ablation tomography (LAT) cross-sectional images of 2-to 3-year-old branches from three ring- (*C. ovata, Q. montana, F. pennsylvanica*; top row) and three diffuse-porous (*P. x canadensis, L. tulipifera, F. sylvatica*; bottom row) trees. B. Raw images were preprocessed for intensity normalization and motion artifact removal. The preprocessed images were binarized (xylem vessels are filled in green). C. Scanning electron microscopy (SEM) images of two adjacent xylem vessels (V) with intervessel pit connections (yellow arrow) of *F. pennsylvanica*. D. Graphical representation of a xylem network generated from 55 slices of images with the centerlines of xylem shown in blue and the intervessel pit connections shown in red. E. Network representations of xylem. (I) Schematic of xylem vessels (blue) that consists of segments (magenta) with endpoints (green) that terminate at the ends of the xylem, at the edge of the image, or at pit connections (orange). Adjacent xylem vessels are connected by intervessel pit connections. (II) Depiction of individual image planes with segmented cross-sections of xylem vessels and intervessel pit connections. (III) Xylem network graphs show xylem connected to neighboring xylems with intervessel pit connections. (IV) Fluid network graphs connect the endpoints of segments to the next segment in the same xylem or to segments in adjacent xylem via pit connections. (V) ϑ-obrazom network representation with segments as nodes.

**Table 1.** Sampling thresholds for laser ablation tomography and binarization analysis.

| **Wood type** | **Tree species** | **Cutting distance between images (μm)** | **Total sample length (mm)** | **Intervessel threshold (μm)** |
| --- | --- | --- | --- | --- |
| Diffuse-porous | *F. sylvatica* | 40 | 70.0 | 2 |
|  | *L. tulipifera* | 50 | 62.9 | 3 |
|  | *P. x canadensis* | 35 | 46.0 | 3 |
| Ring-porous | *C. ovata* | 50 | 98.0 | 4 |
|  | *F. pennsylvanica* | 50 | 271.4 | 5 |
|  | *Q. montana* | 50 | 167.0 | 5 |

We represented 3D structures of xylem using three network schemes that emphasize different aspects (i.e., xylem, fluid, and ϑ-obrazom networks, Fig 1E). The xylem network scheme represents each xylem vessel as a graph vertex (*i-v* in Fig 1E(I)) and the pit connections as the graph edges (Fig 1EII). A xylem vessel can have multiple pit connections to adjacent xylem vessels which are represented as one edge (Fig 1E(III)). The fluid network scheme divides a xylem vessel into pipe-like segments (a-k in Fig 1E(I)) with two endpoints that define the vertices. Some segment endpoints do not have connections because the xylem ends or because it is at the edge of the imaged region. Other endpoints are connected to a segment in an adjacent xylem vessel by an intervessel pit connection where water can move between different xylem vessels or to the next segment in the same xylem vessel (Fig 1E(IV)). The third scheme, the ϑ-obrazom, is the edge-to-vertex dual of the fluid network in which vertices are defined as the fluid network’s segments and two types of edges represent the connection between consecutive segments within the same xylem vessel or pit connections to segments in an adjacent xylem vessel (Fig 1E(V)). While the xylem network is useful for looking at the topology of the 3D xylem, the fluid network and ϑ-obrazom schemes are useful for analyzing the fluid flow and other network characteristics of the xylem.

### Morphological comparisons of xylem vessels and xylem segments

Morphological descriptors in ϑ-obrazom and xylem networks showed similar trends across tree species and wood types with a few exceptions. In the ϑ-obrazom representation, xylem segments of ring-porous tree species had 6-to 14-μm larger diameters than diffuse-porous tree species (Fig 2A.I, Table S 1)). Differences in segment diameter were also detected within the ring-porous tree species, as the segment diameter of *C. ovata* is smaller by 7-8 μm than those of *F. pennsylvanica* and *Q. montana* (Fig 2A.I, Table S 1). As in the ϑ-obrazom networks, the diffuse-porous tree species had similar diameters when averaged across the entire length of the xylem vessel, measured based on xylem networks (Fig 2B.I, Table S 2). However, differences in vessel diameter within the ring-porous tree species ranged between 4 and 11 μm (Fig 2B.I, Table S 2). *F. pennsylvanica* has ~100 μm longer xylem segments than any other tree species (Fig 2A.II, Table S 1), but xylem vessel length from the xylem network does not differ within or across wood types (Fig 2B.II, Table S 2). Many segments do not have any pit connections and the differences in the average number of pit connections per segment do not reveal any consistent patterns between wood types or tree species (Fig 2A.III, Table S 1). *C. ovata* and *F. pennsylvanica* have fewer connections per segment compared to *F. sylvatica* and *P. x canadensis* while *L. tulipifera* and *Q. montana* have fewer connections in comparison to P. x canadensis (Fig 2A.III, Table S 1). In most tree species, the xylem networks showed a xylem vessel is often connected to several other xylem vessels as indicated by the number of connections per vessel, but *C. ovata* stands out for often having many xylem vessels connected to only one or no other xylem vessels (Fig 2B.III, Table S 2). There are no consistent trends in the number of pit connections per xylem vessel across wood types. Finally, *F. pennsylvanica* has 3 to 5 μm longer pit connection length per segment than the other five tree species (Fig 2A.IV, Table S 1), but no differences are detected between the length of pit connections per xylem vessel across tree species (Fig 2B.IV, Table S 2).

**Fig 2.**
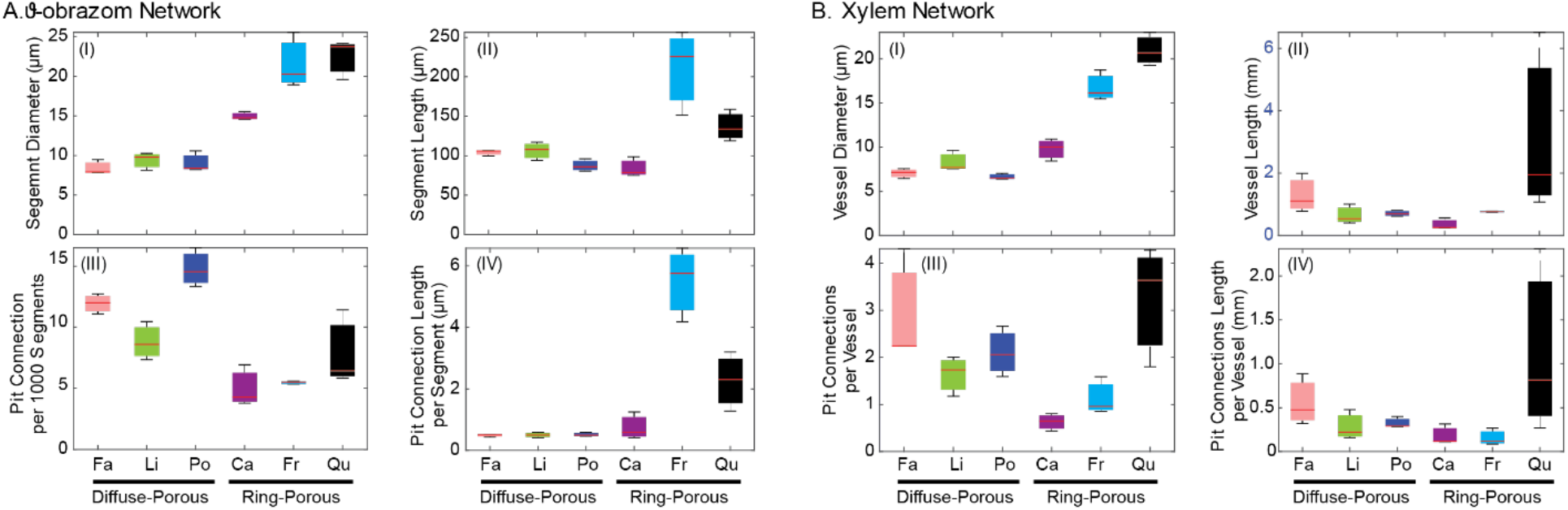
Xylem morphology of three diffuse-porous (*F. sylvatica* (Fa), *L. tulipifera* (Li), *P. x canadensis* (Po)) and three ring-porous tree species (*C. ovata* (Ca), *F. pennsylvanica* (Fr), *Q. montana* (Qu)). A. Morphological characterization of ϑ-obrazom networks. Box plots represent xylem segment diameter (A.I), segment length (A.II), number of intervessel pit connections per 1000 xylem segment (A.III), and lengths of intervessel pit connections measured on segments in ϑ-obrazom (A.IV). B. Morphological characterization of xylem networks. Box plots represent xylem vessel diameter <u>(B.I)</u>, vessel length (B.II), number of intervessel pit connections per xylem vessel (B.III), and lengths of intervessel pit connections measured on vessels in xylem networks (B.IV). Statistical differences are given in Table S 1 and Table S 2.

### Topological comparisons of xylem networks

Using both ϑ-obrazom and xylem networks, we calculated network metrics which can be averaged across samples to represent the species (Fig 3A). Occasionally, a single network metric can predict an interesting property of a network, and a few network metrics suggested segregation between diffuse- and ring-porous wood types. Ring-porous species have a higher ϑ-obrazom density (i.e., ratio of number of edges compared to a fully connected network) compared to diffuse-porous species. The average of ϑ-obrazom network Katz, standard deviation of xylem network clustering, standard deviation of ϑ-obrazom network assortativity, and standard deviation of xylem network assortativity demonstrate similar discrimination between wood types. On the other hand, *C. ovata, P. x canadensis, Q. montana* have higher centrality metrics in terms of edge and vertex betweenness (based on both average and standard deviation) indicating a tendency for some vertices to be more critical than others compared to the other three species in both network representations, showing a trend not related to wood types. The same metrics calculated from the two network representations also suggest contradictory relationships. For instance, based on the xylem network representation, *Q. montana* and *F. sylvatica* have the highest density and eigenvalue compared to all other species, respectively, but not in the ϑ-obrazom networks. There were no obvious patterns that were consistent between different metrics across tree species.

**Fig 3.**
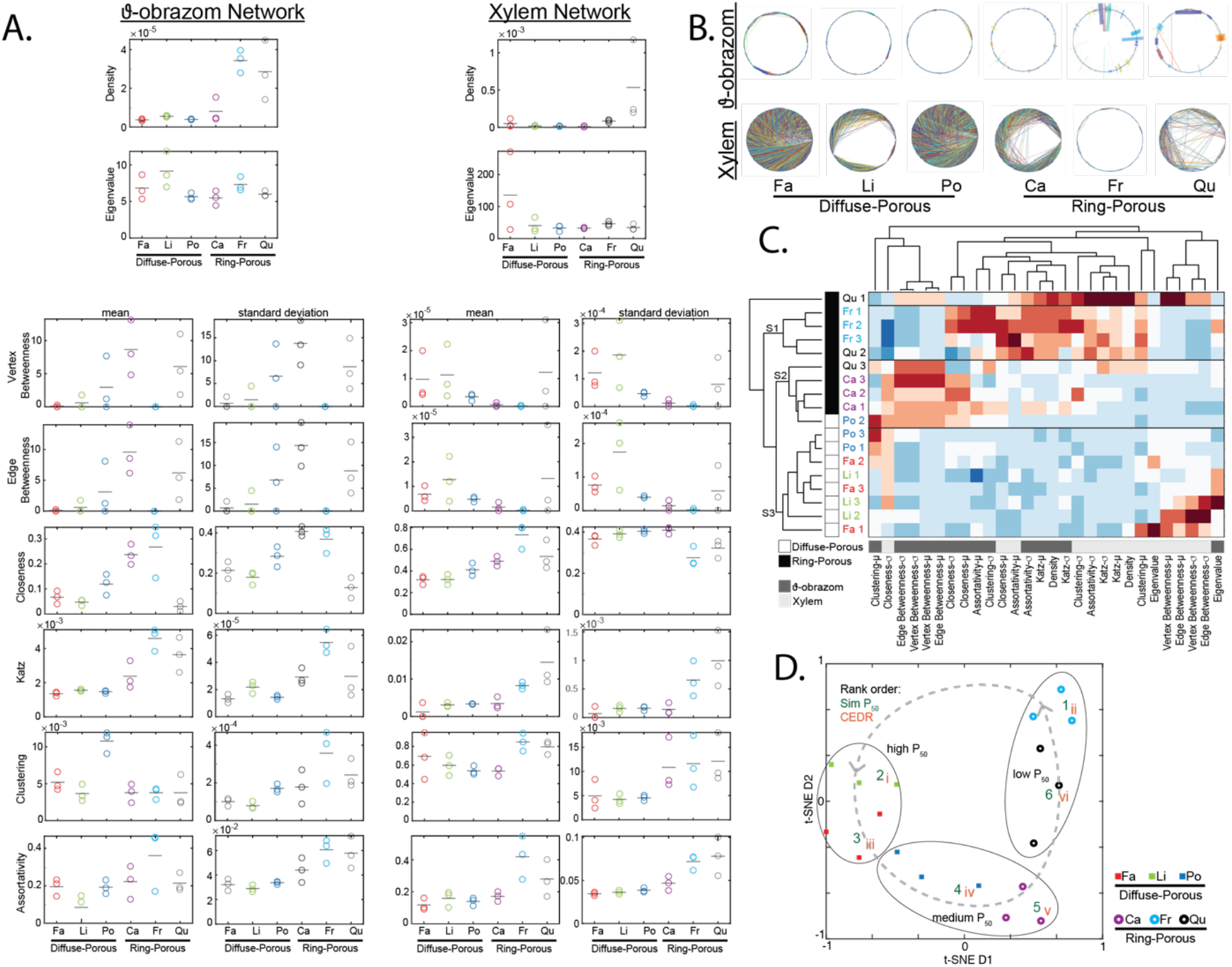
Topological properties of xylem with multivariate analyses. A. Network analysis based on the ϑ-obrazom and xylem networks. Fourteen network metrics (mean (μ) and standard deviation (σ)) from the two network representations of *F. sylvatica* (Fa), *L. tulipifera* (Li), *P. x canadensis* (Po), *C. ovata* (Ca), *F. pennsylvanica* (Fr), and *Q. montana* (Qu). B. Circle graphs from three diffuse- and three ring-porous tree species based on the ϑ-obrazom and xylem networks. All vertices are aligned along the circle perimeter and edges (connections) are drawn between vertices. In ϑ-obrazom networks the edge thickness perpendicular to the circle perimeter is proportional to the total length of the pit connections between two vertices. One circle graph per species is shown as a representative for three replicates per species. For all the graphs see Fig S 5 and Fig S 6. C. Heatmap and hierarchical clustering analysis from both the ϑ-obrazom and xylem networks. Rows represent samples and columns represent the means (μ) and standard deviations (σ) of network metrics shown in B. Metrics based on the ϑ-obrazom network are indicated in dark gray and those based on the xylem network are in light gray. Data in the heatmap are normalized by mean and standard deviation in each column. Each row represents a single sample (numbered) with diffuse- and ring-porous wood types indicated in white and black. D. Dimensional reduction by t-SNE enables visualization of the 28-dimension network analysis (40). Each point represents one sample, with species indicated in colors and wood types distinguished by shape. Distance between points on the 2D graph reflects distance in the full multidimensional space. Numbers on graph indicate the rank order derived from species-averaged simulated P_50_ (Sim P_50_, Arabic numerals in green) and conductance exponential decay rate (CEDR, Roman numerals in red). Ellipses correspond to high, medium, and low values of experimentally measured P_50_ from Fig. 4.

We also used circle graphs (39) to visualize similarities and differences between xylem and 9-obrazom networks across species (Fig 3B). Circle graphs are generated by aligning all the vertices of one network as single points along the circumference of a circle and drawing a line between vertices that are connected. In ϑ-obrazom networks, the thickness of the line indicates length of the pit connection between two connected vertices. The circle graphs of ϑ-obrazom networks show that diffuse-porous tree species have connections between distant conduits, while ring-porous tree species connect more with nearby xylem segments (Fig 3B). The circle graphs of xylem networks suggest that *F. sylvatica and P. x canadensis* are similar, but variability within species is high (Fig S 5 and S 6) and it is difficult to make meaningful comparisons across groups.

### Clustering of multiple network properties as a comparison method

To better relate network properties to functional measures, we chose to combine the topological properties calculated from both xylem and ϑ-obrazom networks as high-dimensional descriptors inspired by bioinformatic techniques for complex data such as genomics (40). These approaches combine measurements from different samples and generate groupings based on multiple metrics with unbiased algorithms. We used a heatmap in which columns represent normalized values of network metrics and each row shows a sample (Fig 3A). Hierarchical Clustering (41) reveals relationships between species using network metrics from both the xylem and ϑ-obrazom network representations. Samples (heatmap rows in Fig 3A) clustered into three main clusters which effectively grouped samples of same tree type, with the exception of one *P. x canadensis* sample (Po 2 in cluster S2) and one *Q. montana* sample (Qu 1) which clustered by itself (first line) (Fig 3C). All samples within the cluster S3 are diffuse-porous tree species and all samples within the cluster S1 are ring-porous species. The cluster S2 is mostly ring-porous species. It should be noted that the single *P. x canadensis* sample that ended up in the S2 cluster is very close to the S3 cluster which contained all the other diffuse-porous samples and all the ring- and diffuse-porous species were grouped together. Network metrics (columns in Fig 3A) were grouped into three main clusters, which contained a mix of metrics from either network representation.

Dimensionality-reduction approaches such as t-SNE (t-distributed stochastic neighbor embedding) (40) are often used to visualize groupings based on similarities across many properties. We used the same data shown in the heatmap (Fig 3C) to generate a t-SNE graph in which each point represents a single sample (Fig 3D). The distance between points can be interpreted as a metric of similarity. Samples from the same species were grouped closely together and the two wood types were clearly separated into clusters (circles and squares in Fig 3D).

### Simulations of network robustness with vessel drop out

The topology and organizational structure of a network determine the impact of loss of network elements (42). Using the fluid network representation we simulated water flow through a sample with a constant pressure difference between the two ends. As a measure of robustness relevant to tree biology, we characterized the change in flow due to the elimination of increasing fractions of randomly selected segments (dropout) for 30 simulations at 1% intervals (Fig 4A and Fig S 4)). Curves were fit to an exponential decay curve parameterized by the conductance exponential decay rate (CEDR, Fig 4A see SI Materials and Methods). In nearly all networks, there was a dropout fraction above which conductance (mmol m^-2^ s^-1^ MPa^-1^) was nearly stopped (Fig 4A). The ring-porous tree species *Q. montana* showed the fastest decrease in relative conductance with a nearly 100% conductance loss at a vessel dropout probability of 2% (Fig 4A). In the diffuse-porous tree species, *F. sylvatica* and *P. x canadensis*, a total conductance loss occurred at about 10% vessel dropout, while the 100% conductance loss of *L. tulipifera* varied between 10% and 40% due to the variability within the samples (Fig 4A). *F. pennsylvanica* had very large variability (Fig 4A). For comparison, similar simulations in blood vessel and synthetic networks are shown in Fig 4B (35). Based on the simulated decrease in relative conductance with increasing vessel dropout (Fig 4A), simulated P_50_ (SP_50_) values were defined as the dropout percentage that resulted in 50% conductance in each sample. A comparison of these values among species revealed that neither the SP_50_ nor the CEDR significantly differ between the species (Fig 4D).

**Fig 4.**
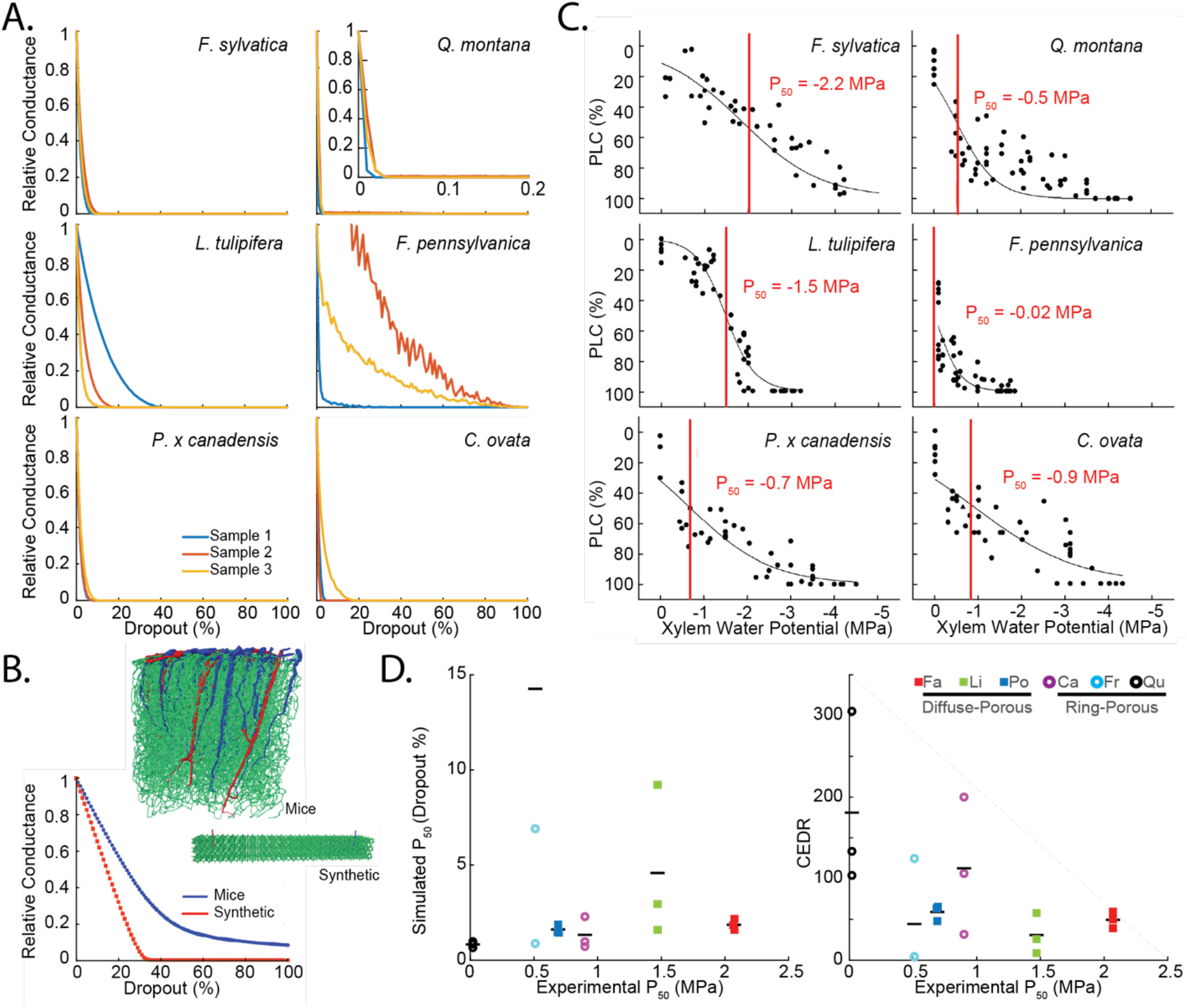
Simulated and experimental fluid dynamic analysis. A. Simulated relative hydraulic conductance as a function of fraction of removed vessel representing embolized vessels (drop out) of three diffuse-porous (*F. sylvatica* (Fa), *L. tulipifera* (Li), *P. x canadensis* (Po)) and three ring-porous tree species (*C. ovata* (Ca), *F. pennsylvanica* (Fr), *Q. montana* (Qu)). Three replicates per species are shown. B. Simulated cerebral blood flow with dropout of capillaries in mouse cortex or in a synthetic network with the same branching order. The relative conductance decreases faster in the artificial network (red) than in the mouse network (blue). Graphs show expansions of the simulations originally published in (35). C. Measured percent loss of hydraulic conductivity (PLC) curves of diffuse- and ring-porous tree species. PLC curves obtained by fitting an exponential sigmoidal function to the data. P_50_ is the xylem water potential at which PLC is reduced to 50% (red line). Values for fit parameters are given in Table S 3. D. Simulated P_50_ values (left) and conductance exponential decay rate (right) versus experimental P_50_. Black line represents mean. The third simulated P_50_ datapoint of *F. pennsylvanica* is 0.35, which is not depicted in the chart. The markers for two lowest simulated P_50_ datapoints and the two highest CEDR datapoints of *P. x canadensis* are overlapped.

### Comparisons between experimental P_50_ values and simulation

We experimentally measured P_50_ values, the pressure at which hydraulic conductance was reduced by 50% from maximum hydraulic conductance from hydraulic vulnerability curves (PLC curves) for each tree species (Fig 4C). P_50_ values typically range from −0.2 to −14.0 MPa-in trees, with more negative values corresponding to greater drought tolerance (22). PLC curves showed that ring-porous tree species had generally higher P_50_ values than diffuse-porous tree species (Fig 4C). However, *P. x canadensis* and *C. ovata* did not strictly follow this trend. *C. ovata* is considered more drought-resistant with a P_50_ value of −0.9 MP, than *P. x canadensis*, which has a P_50_ value of −0.69 MPa. PLC fitting parameters *a* and *b* were all highly significant, except fitting parameter *b* for *F. pennsylvanica*, indicating that a more simplistic model fit would have been sufficient for this species (Table S 3). Neither simulated SP_50_ values nor CEDR showed any strong relationships with experimental P_50_ values, (Fig 4D).

## Discussion and Conclusions

Here, we present for the first time an analysis of xylem vascular networks that characterize the topology and connections of xylem vessels over several centimeter-long branches and the entire width of a year ring. Because the method of reducing vascular networks to graphs can influence the analysis, we used multiple representations to capture the anatomical and morphological features that influence xylem network function (Fig 1D). For example, while the xylem network representation better reported anatomical characteristics of xylem vessels, the ϑ-obrazom network was aimed towards a better understanding of capturing the resistance of water flow through the xylem network.

To evaluate differences between the networks, we took advantage of the large number of network descriptors from graph theory (43) and calculated normalized metrics based on both xylem and ϑ-obrazom networks. We found that functional biological properties or phylogeny were difficult to relate to single or even groups of network properties. Rather than attempt to choose between graphing conventions and network metrics, we combined as much of the information as possible. Inspired by unbiased bioinformatic approaches that identify important groups of properties such as groups of genes that predict a disease better than a single gene, or more recently, complex cell behaviors such as motion (44), we developed a multidimensional approach to characterize xylem networks using 28 different metrics from two different network representations. The combination of different network representations and inclusion of many measures provided a robust way to categorize and compare tree xylem networks. We found that the ring- and diffuse-porous wood types are distinct and separable using an unbiased hierarchal clustering algorithm (Fig 3C). The t-SNE plots provided another useful dimensionality-reduction tool to capture the many aspects of network structures varying across species and environments. Within the t-SNE graphs (Fig. 3D), the individual samples of each species were closely positioned and the two wood types also appeared segregated on the left and right sides of the plot, suggesting that this categorization has biological meaning. Because proximity on t-SNE graphs suggests similarities, trajectories of these graphs can be used to suggest useful relationships. For example, it has been documented that the maturation of different types of immune cells results in trajectories in t-SNE plots that result in changes of gene expression in cells over time (45). The t-SNE plot suggests that although *P. x canadensis and C. ovata* are not the same wood type, the xylem networks are similar. These two species also appeared close together in the hierarchical clustering. While it was difficult to relate the experimental P_50_ to any single network characteristic, species grouped by high, medium, and low experimental P_50_ values on the t-SNE plot along a trajectory (Fig 3D, grey ellipses). Interestingly, we find that the rank order of species in increasing simulated P_50_ values (numbers, Fig 3D) and CEDR followed the same trajectory, with the exclusion of *F. pennsylvanica. F. pennsylvanica* CEDR estimations showed some inconsistencies that could be related to sample imaging difficulties. Either excluding *F. pennsylvanica* or starting the t-SNE trajectory with *F. pennsylvanica*, a t-SNE trajectory can be drawn that recapitulates the rank order of both the simulated P_50_ and CEDR values. Because there are many methods for graphically representing high-dimensional data sets and the final distributions depend on algorithm parameters, these trajectories may vary depending on these choices. However, they are a subjective tool to compare data and predict groups that may share some similar properties in experiments or in the field (46). The hierarchical clustering and t-SNE analysis are unbiased analyses, which can accommodate many more types of measurements and samples. Future studies could use such multidimensional techniques to better compare differences between species or environmental conditions.

We used computational fluid dynamic methods combined with simulation of stochastic appearance of embolisms by random dropout of network elements, to generate a functional metric of robustness that relates xylem structure and intervessel connections. In addition to comparisons between tree species, this simulation allows comparisons to other types of networks. Simulated flow in networks of blood vessels from the brains of mice and humans was maintained at approximately 50% when 25% of the capillaries were occluded (35), which indicates a much higher robustness of flow to occlusions than in the plant vascular systems (Fig. 4C). A synthetic network with the same number of vessels per branch as the blood vessel network (47) also showed more robustness to occlusions than the trees, but less than blood vessels. This suggests that robustness to sporadic occlusion is of relatively low importance in tree vasculature. A possible interpretation is that although intervessel connections between xylem vessels can confer robustness of flow against loss of xylem vessels by providing redundant pathways for water transport, these connections can also threaten water transport efficiency during drought stress periods so that there is evolutionary pressure to limit such connectivity (9,32). Under high tension, embolisms can spread across pits into adjoining xylem conduits by aspiration, further preventing water transport. These two contrary arguments highlight the necessity for characterizing the architecture of xylem networks to identify network traits affecting tree survival.

While our flow simulations provide a valuable tool to compare networks, they only mimic conductance change resulting from randomly occurring embolism events most relevant for flow under low tension (18, 48). Our model does not include the effects of embolisms propagating from air aspiration through intervessel pit membranes, e.g., air-seeding (3), and therefore, currently, we are unable to conclude if higher connectivity leads to more embolism-resistant or more vulnerable networks. Furthermore, in our fluid simulation, we model water transport with a constant pressure gradient across the network and do not include equations that connect the local pressure in the xylem to the xylem drop-out procedure in the analysis. As a result, we only represent the general vulnerability of xylem networks to embolism events and cannot make any predictions about the tension threshold at which the rapid conductance decline occurs. Thus, even though our model predicts a severe conductance decline with increasing vessel dropout rate, cavitation might not occur until severe drought stress conditions (severe xylem tension). In order to simulate air-seeding in xylem networks, a follow-up simulation must be performed in which randomly dropped vessels will also trigger the removal of all connected vessels.

Combinations of morphological and topological differences based on both graph representations reveal the complexity of the correlation between the xylem network characteristics and drought vulnerability. In this study, we were able, for the first time, to produce a large 3D, image data set and to reconstruct xylem networks based on actual anatomy. Furthermore, we presented different metrics to characterize network topology and a bioinformatics-inspired method to combine all of these metrics to serve as a comparative framework for studying xylem network robustness and other complex network properties.

## Materials and Methods

For additional details on materials and methods see SI Materials and Methods.

### Plant material

Two two-year-old branches of three individual trees of either ring-porous (*Quercus montana, Fraxinus excelsior, Carya ovata*) or diffuse-porous (*Fagus sylvatica, Populus x canadensis, Liriodendron tulipifera*) species were harvested respectively for determining vessel length distribution, measuring P_50_ values, and segmenting xylem networks between April 2016 and June 2017. Selected mature tree species were grown on Cornell University campus, and tree replicates were chosen based on their proximity to each other (Ithaca, NY; lat. 42.44° N, long. 76.44° W). Ithaca has a moderate continental climate with an average annual temperature of 8.5°C and an average annual rainfall of 982 mm (49). Branches were harvested from the lower canopy (~3 m above ground) with a pole pruner. For performing PLC curves (see supporting information) and laser ablation tomography, branches (10-20 cm long) were harvested during the day, while for hydraulic conductance measurement branches (60-100 cm long) were collected around midnight to prevent artificial embolism occurrence in the xylem conduits (50). After harvesting, branches were brought to the lab. For anatomical measurements and hydraulic conductance measurements, samples were processed on either the same or the next day (please see respective section). Samples for laser ablation tomography were placed in a sealed plastic bag and shipped to L4iS (State College, PA), where samples were refrigerated until processing.

### Anatomical measurements

#### Sample length

Sample length for xylem binarization was determined from the vessel length distribution, which was calculated for each species based on the silicon injection technique (16, 29). The cutting distance for the branches of each species was set to the 75^th^ percentile of the vessel class length for diffuse-porous tree species and to the 50^th^ percentile for ring-porous tree species (Table 1). These percentile thresholds were set differently for wood types to ensure adequate sampling while minimizing the number of images acquired.

#### Intervessel wall thickness

Scanning electron microscopy (SEM, Zeiss 1550, Oberkochen, Germany) was used to measure intervessel wall thickness. The distance between two adjacent vessels was determined as the minimum distance at which two vessels were connected (Fig 1C). Three branches per individual tree (nine branches per tree species) were harvested and cut into 5 mm samples with a sliding microtome (American Optical, 680 sliding microtome, Spencer Lens Co, Buffalo, NY). Then, samples were dehydrated in a series of 25%, 50%, 70%, 95%, and 100% ethanol, and dried at room temperature. Samples were coated with gold-palladium for 20 seconds at a current of 20 mA and imaged at a voltage of 3.0 kV and a current of 0.21 nA. The intervessel wall thickness was measured in ImageJ (51) based on a minimum of 60 adjacent vessels.

### Image processing pipeline

Image processing and analysis steps for extracting the three-dimensional structure of xylem networks and for understanding the fluid dynamics through these networks are described in the following paragraphs with further details in supplementary materials.

#### Image acquisition

Laser ablation tomography (LATscan, Laser for Innovative Solutions, State College, PA, USA) was used to generate three-dimensional images by acquiring digital cross-sectional images of the samples, (Fig 1A). A coherent Avia 355-7000 Q-switched ultraviolet laser (Coherent, Santa Clara, CA, USA) with a pulse repetition rate of 25 kHz and wavelength of 355 nm was used for ablating the samples. The pulse duration of the laser was less than 30 ns and supplied pulse energy of approximately 200 mJ. The galvanometer used to scan the laser beam to make the ablation plane was a Scanlab HurryScan 10 (Scanlab, Siemensstr 2a, 82178 Puchheim, Germany). Samples were fixed to a cantilever and connected to the mechanical stage along its travel axis, then fed into the ablation plane using an Aerotech linear drive stage (Aerotech, Inc., 101 Zeta Dr, Pittsburgh, PA 15238, USA), with the cutting distance ranging from 35 μm to 50 μm. Images were captured via a Canon 70D camera equipped with a Canon Macro Photo Lens MP-E 65 mm 1:2.8 1-5X. The images captures were 5472 x 3648 pixels at a resolution of 1 μm per pixel.

#### Xylem vessel image processing and network graph generation

After motion artifacts were corrected (see supplements), a trained scientist manually segmented xylem vessels images with at least 100 vessels in three samples from each species with at least 31 consecutive images were manually annotated as the ground truth (36). Due to the significant anatomical differences between vessel morphologies across tree species, species-based models outperformed the universal tracing model.

Since each sample data set contained between 1000 and 5000 images, 500-image sub-stacks were stored in a HDF5 binary data format to be segmented using the CNN model. After this initial binarization task, the results from sub-stacks were concatenated to form the complete binarization data set for each sample. To remove small artifacts, we applied a dilation morphological image filter with a disk kernel of radius 1 voxel to remove the boundary of vessels followed by a 3D median filter with a 3-voxel box kernel to smooth the vessel boundary and fill the holes within the vessels.

#### Fluid dynamics simulations

The pressure drops within a 3D xylem segment can be modeled based on the Hagen–Poiseuille law by considering xylem vessels as circular cross-sectional pipes and the resistance of the intervessel connections as infinitely thin plates with perfectly circular pores with resistance as a function of the equivalent pore size and the number of pores in the intervessel connection. The hydraulic network based on the xylem segments and intervessel connections system was modeled in a linear system of equations.

## Statistics

Statistical analysis was performed in JMP Pro 14.0.0 (SAS Institute Inc., Cary, NC.) or Matlab (MathWorks, Natick, MA). All tests were performed with probability level *P* < 0.05. Differences between tree pieces were calculated using ANOVA, and multi-comparison corrections were done using the Tukey-Kramer method. For calculating differences in intervessel wall thickness among tree types, the dependent variable was log transformed to fulfill model assumptions.

## Data Availability

All networks in Matlab format are available from a publicly-accessible, archival website by Cornell University https://ecommons.cornell.edu along with example images.

## Acknowledgments

Authors would like to thank Peter Doerschuk, Naoki Masuda, and Chris Schaffer for their constructive feedback on this manuscript, and Benjamin Hall for his work on LATscan.

## Supplementary Information Appendix

### SI Materials and Methods

#### Anatomical measurements

##### Sample length

For the vessel injection technique, six branches (each ~60 cm in length) per species were cut, brought into the lab, and flushed for 1 hour at 70 kPa with a 20 mM KCL solution to remove native embolisms (52). Then, basal ends of the branches were connected via silicon tubing to a nitrogen gas tank and injected with a 10:1 two-component silicone elastomer (RTV141 A&B, distributed by Hisco, Somerset, NJ, USA) at 70 kPa overnight. Prior to injection, the silicon mixture was degassed under vacuum and infused with a UV stain that was dissolved in chloroform (Ciba Uvitex OB, Ciba Specialty Chemicals, Tarrytown, NY) in order to separate silicone injected xylem vessels from empty vessels for imaging analysis. After the silicon cured (~2 days), branches were sectioned at six cutting distances from the injection site with a sliding microtome (American Optical, 680 sliding microtome, Spencer Lens Co., Buffalo, NY). The respective cutting distances (L_i_) were determined with the following equation

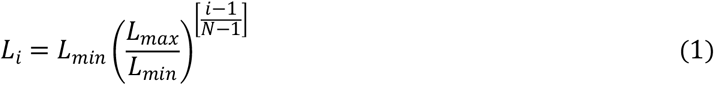

with L_min_ equal to 0.5 cm beyond the injection point, and L_max_ the distance at which 2% of the vessels were detected under the fluorescence microscope, and N the total number of cuts. Then, cross sections were mounted in glycerol and magnified with a 20x objective and imaged with a camera (Retiga Exi CCD camera, QImaging, Burnaby, BC, Canada attached to a fluorescence microscope (Olympus BX50, Olympus Scientific Solutions, Waltham, MA, US). Fluorescent silicon injected vessels from the most recent formed year ring were counted and averaged over species. Last, the vessel length distribution and the average vessel length was calculated for each species on the basis of equations reported in previous studies (53, 54) with the objective to fit the silicon-injected vessel counts with a Weibull function and to use the best fit to calculate the second derivate, from which the vessel length distribution was calculated (Fig S 1).

#### Image processing pipeline

##### Motion artefact compensation

Motion artifact is one of the main challenges for images processing of 3D vasculature network images. Apparent motion can be caused by any combination of gradual shift due to sample curvatures, sudden shift, or rotation due to sample realignment, poor focus, and burned crosssection. To compensate for these different motion artifacts, two different methods were utilized. For each method, the hyperparameters were optimized based on the raw images and type of motion artifacts detected in the analysis. Furthermore, the computational complexity and scalability also had to be considered in the motion artifact compensation, given that the 3D images could contain up to 5000 slices. Since there were no local artifacts due to distortion and motion in the images, a rigid registration (i.e., translation and rotation) was sufficient. Registration methods used a similarity metric as an input to the cost function and optimization of the cost function to find the registration parameters.

The first method that was applied to correct motion artifacts used mean square error *(MSE)* (Eq. 2), where N and M are numbers of rows and columns in the image and I and I’ are the intensity of a particular pixel in two images, as the similarity metric and the regular step gradient descent optimization, which follows the gradient of the cost function toward the direction of extrema with reducing the step function when the gradient changes direction.

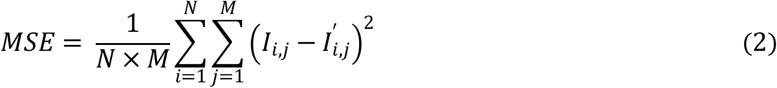

A random grid search was used to find the optimal hyperparameters for the optimizer and detected found two optimal sets of hyperparameters. The ensemble of two optimizers based on the optimal sets of hyperparameters was able to handle most of the cases, and it was successful when applied to the test cases with small sample sizes. The failure modes include uneven illuminations, laser-burned cross-sections, color-distorted images, and sudden dramatic changes. On the other hand, even after multi-thread parallelization, this method has a long run time, in addition to the need for required manual treatments of the remaining failure cases.

We devised the second method based on the work by Evangelidis and Psarakis (55) to overcome the drawbacks of the first method specifically for samples with large sample sizes. They adopted the enhanced correlation coefficient (56) as the similarity measure. This measure has preferable characteristics such as being invariant to contrast and brightness differences (i.e., photometric distortions) as well as having a corresponding linear approximation expression with a closed-form solution, which facilitates the optimization of the original non-linear measure. Additionally, Evangelidis and Psarakis proposed an iterative gradient based searching function (e.g., forward additive refinement algorithm) to optimize the original non-linear measure using the linear approximation (55). In order to tackle multi-scale motion artifacts, the transformation parameters were estimated in a pyramid fashion using 10x scaled-down versions of the two images and then fine-tuned the parameters using the original resolutions. Since the images did not suffer from local motion artifacts, only the estimation of the global transformation parameters within two consecutive images was required.

In order to parallelize the process, images were divided into groups of 10 consecutive images and registered independently based on the first image in the group. Then, sequentially starting from the second image group, the first image of each image group was aligned to the last image of the preceding image group and the rest of the images within the group were warped using the same transformation. Finally, a failure detector was implemented by using the mean structural similarity index (57) between two consecutive images. Within every 11 consecutive pairs of images, the middle pair is considered as a failure case if its similarity index is at least 5% lower than the median of the 11 similarity indexes. The failure cases were registered again using the same algorithm (or in rare cases manually if the algorithm failed repeatedly) followed by warping all the following images using the same transformation parameters. This process was repeated until no failure cases were detected.

##### Preliminary image processing

We developed a graphical user interface (GUI) in Matlab for image processing and network analysis to facilitate the preliminary study analysis and the study design (e.g., cutting distance determinations and methodological feasibility risk analysis). The GUI allows the user to load a 3D stack of images and select the region of interest in addition to tuning the image processing parameters. The image processing pipeline started with the application of a top hat filter followed by a bottom hat filter. Top hat filter subtracts the morphological opening of the image from the original image and similarly bottom hat filter subtract the original image from the morphological closing of the image. The morphological opening consists of an erosion of image followed by dilation, and in contrast, morphological closing consists of a dilation followed by an erosion. The erosion and dilation are the two basic mathematical morphology operations, which require a structuring element (i.e., kernel) to operate. The combination of top hat and bottom hat filters treat small objects with very high or low intensity. Next, we sharpened the image by subtracting the smoothed image (using a Gaussian lowpass filter multiplied by a constant) from the original image. Next step the grayscale image is binarized using an adaptive threshold defined using Otsu’s method (58) in addition to ensuring the preservation of the boundaries between adjacent vessels. Finally, the small isolated objects were removed, and the holes were filled. The vessel cross-section centroids were identified to produce the graph representation of vasculature networks using the binarization results. The GUI save the binarization, graph representation, and the adjacency matrix of vessel connections to the output data file. The manually annotated xylem vessels were used as ground truth and later used for training DeepVess to develop tree species specific xylem binarization models (see below).

##### Imaging interval

The maximum sample length was restricted by a combination of technical limitation and sample trait. The digital camera tended to overheat with increasing sample length due to the amount of image taken. Additionally, the degree of natural bending of the branches increased with sample length. Consequently, samples tended to move out of the imaging plane with increasing sample length and we were unable to realign the sample with imaging plane by simply pushing the sample back in the imaging plane. To ensure image quality we chose maximum sample length to be the 75^th^ percentile of vessel class length, that was determined by vessel length distribution, for diffuse-porous tree species and 50^th^ percentile for ring-porous tree species (Fig S 1, Table S 4). To determine the maximum distance between images, a preliminary experiment was performed in which 31 slices of a branch sample of all six tree species were ablated with a cutting distance of 5 μm (total length of 150 μm). Then, a minimum of 100 xylem vessels per sample were selected and manually traced in all 31 slices using ImageJ (51) while following a previously published manual binarization protocol (36). We then measured the binarization similarity between the first slice and the following slices based on the ratio of the intersection area of two adjacent vessel cross-sections and the first slice vessel cross-section area averaged over all detected vessels to determine the maximum distance between two images such that we observed at least 50% averaged cross-sectional overlaps with an upper limit of 50 μm. We found this distance to be 50 μm for all ring-porous tree species, 40 μm for *F. sylvatica*, 35 μm for *P. x canadensis*, and 50 μm for *L. tulipifera*. These cutting distances were then used for all further analysis (Table 1).

##### Xylem graph generation

We utilized the xylem image binarization results to generate the graph representation of xylem networks to study and characterize them using network analysis and fluid mechanics. Since 3D xylem vessels extend the length of the branch, xylems appear in cross-section in each image slice. Cross-sections were defined from the segmented image slices as regions of connected voxels. Xylem vessels were connected by two types of connections. First, junctions were defined where xylem vessels merged and vessel wall is invisible. Junctions were defined whenever two cross-sections overlapped for two or more sequential images. Xylem segments were defined to start and end with a junction. Second, because intervessel connections consisting of pits between xylem were not directly visible in the images, they were assumed to occur in regions at which the thickness of the wall between two xylem vessels dropped below a threshold value. The thresholds for each species were determined from intervessel thicknesses from SEM cryo-image anatomical analysis and were set at the 95^th^ percentile of measurements (Fig 1C).

In order to characterize these networks using morphological measures, each cross-section was fit to an ellipse, and the shortest axis was defined as the diameter. The diameter of each xylem vessel was defined as the median of the measured cross-sections. Correspondingly, the xylem segment length, number of intervessel connections, and total length of intervessel connections are measured for each xylem segment. Morphological measures were calculated and represented for the ϑ-obrazom and xylem networks. Fluid networks were used to simulate fluid dynamics in xylem networks in response to embolism events (see “fluid dynamics in xylem networks” section). The ϑ-obrazom of fluid networks were studied for topology.

#### Empirical conductivity measurements

Vulnerability curves were performed using the bench-top dry-down method (Tyree and Dixon, 1986). Around 60 cm long branches were cut around midnight, immediately double bagged, brought to the lab, and the cut end was put into water. The following morning, the branches were spread out on the bench-top, single leaves were bagged for allowing leaf water potential (Ψ_leaf_) to equilibrate with the branch water potential (Ψ_branch_) and dried down for varying amounts of time to archive a range of different (Ψ_leaf_). During this timeframe, Ψ_leaf_ was taken on the bagged leaves with a water status console (Soilmoisture Equipment Corp., Goleta, CA). After reaching the desired Ψ_leaf_, branches were double bagged, equilibrated for 12 hours, and remeasured. Then, branches were cut under water to 10 cm long segments and inserted into a custom-built low-pressure flow meter (Melcher *et al*., 2012) by attaching one end of the branch segment to a reservoir that was filled with a (0.1 μm) filtered 20 mM KCl of perfusion solution, and the other end of the branch segment to an analytical balance (HR-200, A&D, Elk Grove, Il). Then, the initial flow rate (Q) was measured. The hydraulic pressure difference between sample and solution reservoir was kept constant between 1.5 kPa and 3.0 kPa during the measurements depending on the tree species. Afterward, branch segments were flushed for 1 hour at 100 kPa with the perfusion solution, and the max flow rate was measured by reinserting the stem segments into the low-pressure flow meter. The unit-length hydraulic conductivity (K) was determined by

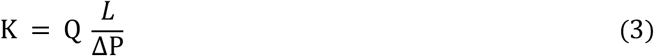

where, L is the length of the sample, and ΔP is the hydraulic pressure gradient. After the measurements, the xylem cross-sectional area was determined with a caliper and the specific hydraulic conductivity determined by dividing K by the cross-sectional areas. Lastly, the percent of hydraulic loss of conductivity (PLC) was calculated by

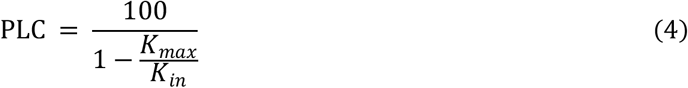

where, K_max_ is the maximum specific conductivity after flushing, and K_in_ is the initial specific hydraulic conductivity. The PLC data were fitted with an exponential sigmoidal equation of

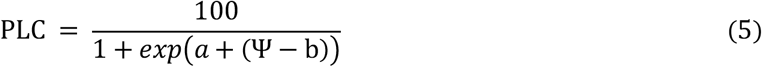

where, a and b are fitting parameters, whereby a describes the slope of the curve and b represents the position of the curve on the x-axis at 50% PLC (59). The significance levels of the parameters were calculated based on this fit.

#### Embolism simulations

The pressure drops within a 3D xylem segment can be modeled based on the Hagen–Poiseuille law by approximating xylem vessels as circular cross-sectional pipes and by assuming that water, which is a incompressible and Newtonian fluid, flows in a creeping regime effectively correlating the fluid flow rate within the pipe (Q) with the pressure drop (ΔP) as formulated in Eq. 6. Hence, the segment resistance is defined as Eq. 7, where *μ* is the dynamic viscosity of water at 25°C, *D* is the xylem segment diameter, and L is the xylem segment length (Loepfe*et al*., 2007).

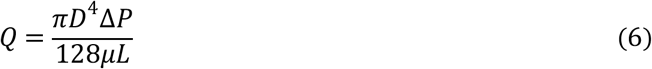

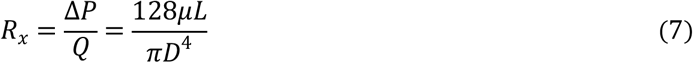

Sperry and Hacke (60) modeled the resistance of the intervessel connections (*R*_i_) as infinitely thin plates with perfectly circular pores with resistance as a function of the equivalent pore size (*D_e_*) and the number of pores in the intervessel connection (*n_p_*) defined in Eq. 8. Since we can assume that n_p_ is proportional to the intervessel connection length (*L*_i_ in Eq. 9), which are measured based on 3D images of samples, we modeled the intervessel connection resistance (*R*_i_) as a function of *L*_i_, which is a proxy for the *n_p_* (Eq. 9). For vascular networks the same approach is used, except that inter vessel resistance is zero and blood taken as a non-Newtonian fluid (35).

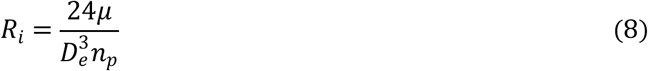

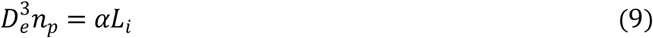

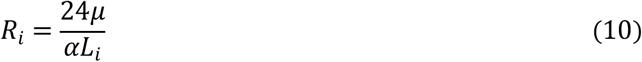

The hydraulic network based on the xylem segments and intervessel connections was modeled using Eq. 7 and Eq. 10 where α was considered as one. The system was then represented in a linear system of equations, and the solution was acquired using one of the sparse systems of linear equation solver methods (e.g., Cholesky solver) depending on the characteristics of the linear systems. Since the problem is linear and the relative variation is needed, a unit pressure difference was applied between the two ends of the longest connected segment within each tree sample, and the flow within the sample was measured to calculate the sample conductance (*C=Q/ΔP*). Thirty simulations were conducted for each dropout probability (*DP*) ranging from 0% to 100% in order to simulate different embolism events. The relative conductance (C’=C/C_max_), which is the ratio of conductance to the baseline conductance at 0% dropout was reported for each sample. Embolism simulations were calculated based on xylem binarization results. However, due to image qualities, fluid simulations could not be performed in all samples on the entire length of the segmented image stack (Table S4). The relative conductance curves were fit to an exponential decay curve parameterized by the conductance exponential decay rate (CEDR).

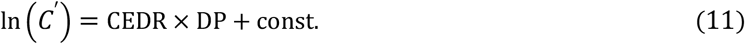

For blood vessel networks the same approach was used, except that intervessel resistance is zero and blood was modeled as a non-Newtonian fluid (35).

#### Network topological analysis

The topological metrics characterize the graphs in terms of the relationships within the vertices. Topological metrics are measured on the graph as whole (e.g. density) or measured for each vertex or edge independently (i.e. closeness). The edge- or vertex-based topological metrics results in measurement distributions, which can be summarized in terms of their mean (*μ*) and standard deviation (*σ*). The implications of the topological metrics can be illustrated using the analogy based on the United States Highway System (USHS).

The *density* measures the degree of connectedness within the network based on the ratio of the current number of edges and the maximum possible number of edges. In USHS, the number of highways in the current system compared to the case that each city is connected to all other cities directly.

The *centrality* metrics, which are topological metrics measuring the importance of edges, vertices, or paths. The centrality metrics utilize the identified shortest path between all pairs of vertices. The *edge*- and *vertex-betweenness* measure number of shortest paths that include an edge or vertex, respectively. Similarly, *closeness* measures the average shortest path to other vertices demonstrating the level of influence of this vertex on other vertices. In USHS, the betweenness illustrate the level of effects in the case of city entrances or a highway closure due to constructions or catastrophic events. The betweenness is the number of shortest paths that are going to be affected due to the closure. Correspondingly, closeness shows the important of a city and how it’s congestions may have a ripple effect on other cities.

The *clustering* and *assortativity* measure the amount of the closed loop within the system and the level of clustering with similar edge types. In USHS, when a city is connected to two different cities, whether those two cities are connected is correlated with the level of interconnections and clustering in the network. The *assortativity*, measures how similar highways are connected for instance main highways vs. controlled access highways (e.g., US I-95 vs. US I-495).

The graph connection can be recoded using the *adjacency matrix*, whose indices represents the vertices, and the entry are non-zero when there is an edge between the corresponding vertices. The largest *eigenvalue* and its eigenvector of the adjacency matrices represents the importance of the main pattern of edge connections within the graph and *Katz* utilized the same concept to measure the relative degree of influence of vertices.

## SI Tables and Figures

**Table S 1.**
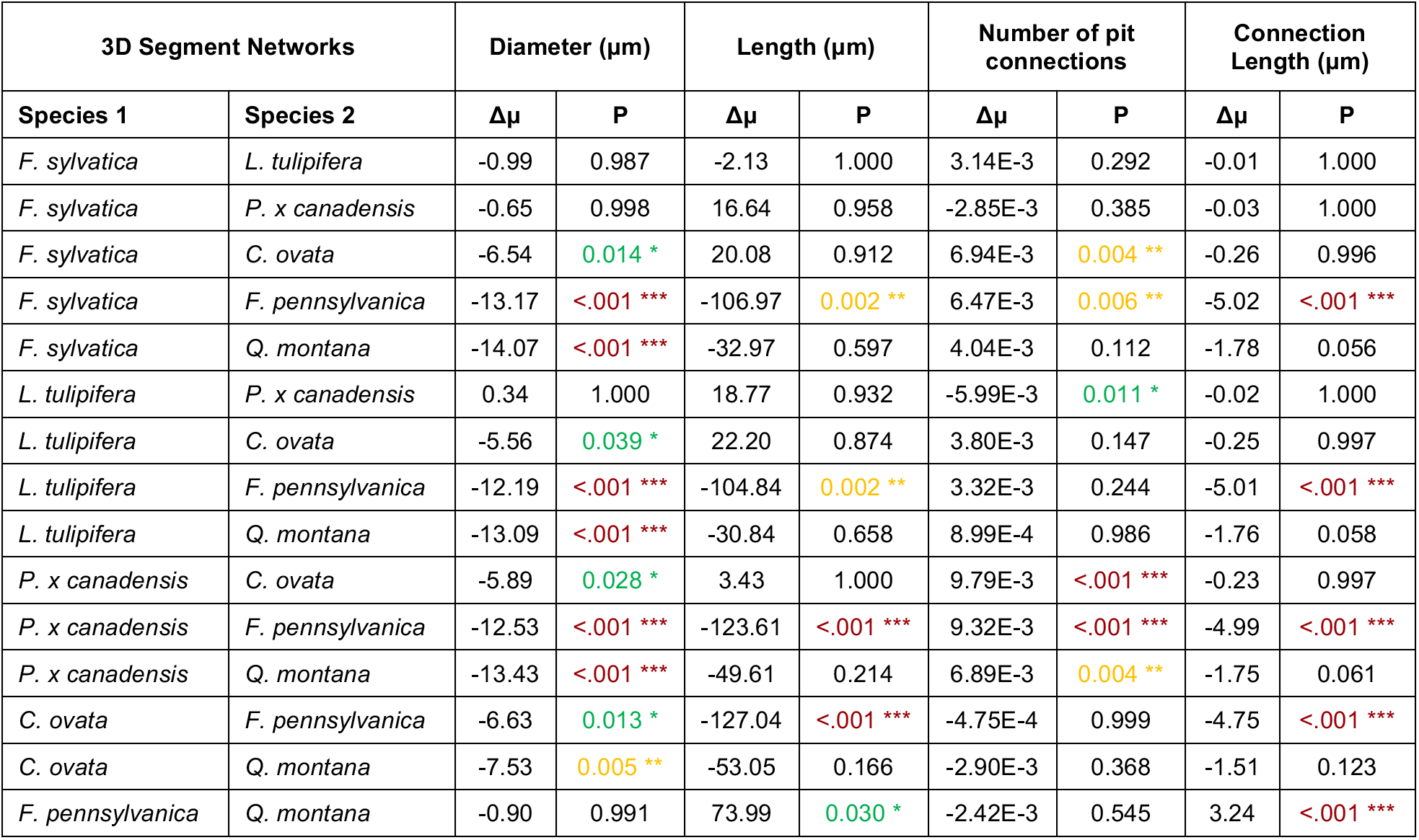
Comparisons between means of vessel diameter, vessel length, intervessel connection frequency and connection length of three ring- (*F. sylvatica, P. x canadensis, L. tuilipifera*) and three diffuse-porous tree species (*C. ovata, Q. montana, F. pennsylvanica*) based on the ϑ-obrazom analysis (ANOVA and Tukey-Kramer multi-comparison correction).

**Table S 2.** Comparisons between means of diameter, length, intervessel connection frequency and connection length of three ring- (*F. sylvatica, P. x canadensis, L. tuilipifera*) and three diffuse-porous tree species (*C. ovata, Q. montana, F. pennsylvanica*) based on the xylem network analysis (ANOVA and Tukey-Kramer multi-comparison correction).

| Xylem Networks |  | Diameter (μm) |  | Length (μm) |  | number of pit connections |  | Connection length (μm) |  |
| --- | --- | --- | --- | --- | --- | --- | --- | --- | --- |
| Species 1 | Species 2 | Δμ | P | Δμ | P | Δμ | P | Δμ | P |
| *F. sylvatica* | *L. tulipifera* | -1.25 | 0.833 | 642.79 | 0.985 | 1.30 | 0.391 | 275.25 | 0.973 |
| *F. sylvatica* | *P. x canadensis* | 0.42 | 0.998 | 586.31 | 0.990 | 0.83 | 0.789 | 235.66 | 0.986 |
| *F. sylvatica* | *C. ovata* | -2.74 | 0.169 | 941.26 | 0.929 | 2.30 | 0.035 * | 379.40 | 0.905 |
| *F. sylvatica* | *F. pennsylvanica* | -9.73 | <.001 *** | 526.58 | 0.994 | 1.80 | 0.127 | 402.89 | 0.882 |
| *F. sylvatica* | *Q. montana* | -13.93 | <.001 *** | -1885.4 | 0.458 | -0.31 | 0.996 | -571.80 | 0.655 |
| *L. tulipifera* | *P. x canadensis* | 1.67 | 0.618 | -56.48 | 1.000 | -0.47 | 0.974 | -39.59 | 1.000 |
| *L. tulipifera* | *C. ovata* | -1.49 | 0.717 | 298.47 | 1.000 | 1.01 | 0.634 | 104.15 | 1.000 |
| *L. tulipifera* | *F. pennsylvanica* | -8.48 | <.001 *** | -116.2 | 1.000 | 0.50 | 0.966 | 127.64 | 0.999 |
| *L. tulipifera* | *Q. montana* | -12.68 | <.001 *** | -2528.2 | 0.193 | -1.60 | 0.203 | -847.05 | 0.280 |
| *P. x canadensis* | *C. ovata* | -3.16 | 0.089 | 354.95 | 0.999 | 1.48 | 0.268 | 143.74 | 0.999 |
| *P. x canadensis* | *F. pennsylvanica* | -10.15 | <.001 *** | -59.73 | 1.000 | 0.97 | 0.667 | 167.23 | 0.997 |
| *P. x canadensis* | *Q. montana* | -14.35 | <.001 *** | -2471.7 | 0.210 | -1.13 | 0.523 | -807.46 | 0.323 |
| *C. ovata* | *F. pennsylvanica* | -6.99 | <.001 *** | -414.68 | 0.998 | -0.51 | 0.965 | 23.49 | 1.000 |
| *C. ovata* | *Q. montana* | -11.19 | <.001 *** | -2826.6 | 0.122 | -2.61 | 0.016 * | -951.19 | 0.187 |
| *F. pennsylvanica* | *Q. montana* | -4.20 | 0.017 * | -2411.9 | 0.229 | -2.10 | 0.059 | -974.69 | 0.170 |

**Table S 3.** Values of coefficients a and b from Eq. 5, where a describes the slope of vulnerability curves presented in Fig S 1 is the predicted water potential at which 50% loss in hydraulic conductance occur. Significant parameters are marked by asterisks (**** = P < 0.0001; * = P < 0.05).

| Tree species | a | b |
| --- | --- | --- |
| *F. sylvatica* | 0.84*** | -2.07**** |
| *L. tulipifera* | 2.92*** | -1.47**** |
| *P. canadensis* | 1.07*** | 0.69**** |
| *Q. montana* | 1.13*** | -0.51**** |
| *F. pennsylvanica* | 3.64*** | -0.02* |
| *C. ovata* | 0.85*** | -0.90*** |

**Table S 4.** Image stack length used for embolism simulation model for each species and replicate.

| Tree species | replicate | Analyzed sample length (mm) | Total sample length (mm) |
| --- | --- | --- | --- |
| *F. sylvatica* | 1 | 70.0 | 70.0 |
|  | 2 | 70.0 | 70.0 |
|  | 3 | 70.0 | 70.0 |
| *L. tulipifera* | 1 | 27.8 | 62.9 |
|  | 2 | 62.9 | 62.9 |
|  | 3 | 62.9 | 62.9 |
| *P. x canadensis* | 1 | 46.0 | 46.0 |
|  | 2 | 46.0 | 46.0 |
|  | 3 | 46.0 | 46.0 |
| *C. ovata* | 1 | 97.8 | 98.0 |
|  | 2 | 97.8 | 98.0 |
|  | 3 | 97.8 | 98.0 |
| *F. pennsylvanica* | 1 | 17.8 | 271.1 |
|  | 2 | 70.9 | 271.1 |
|  | 3 | 77.5 | 271.1 |
| *Q. arbor* | 1 | 160.3 | 167.0 |
|  | 2 | 153.7 | 167.0 |
|  | 3 | 158.7 | 167.0 |

**Fig S 1.**
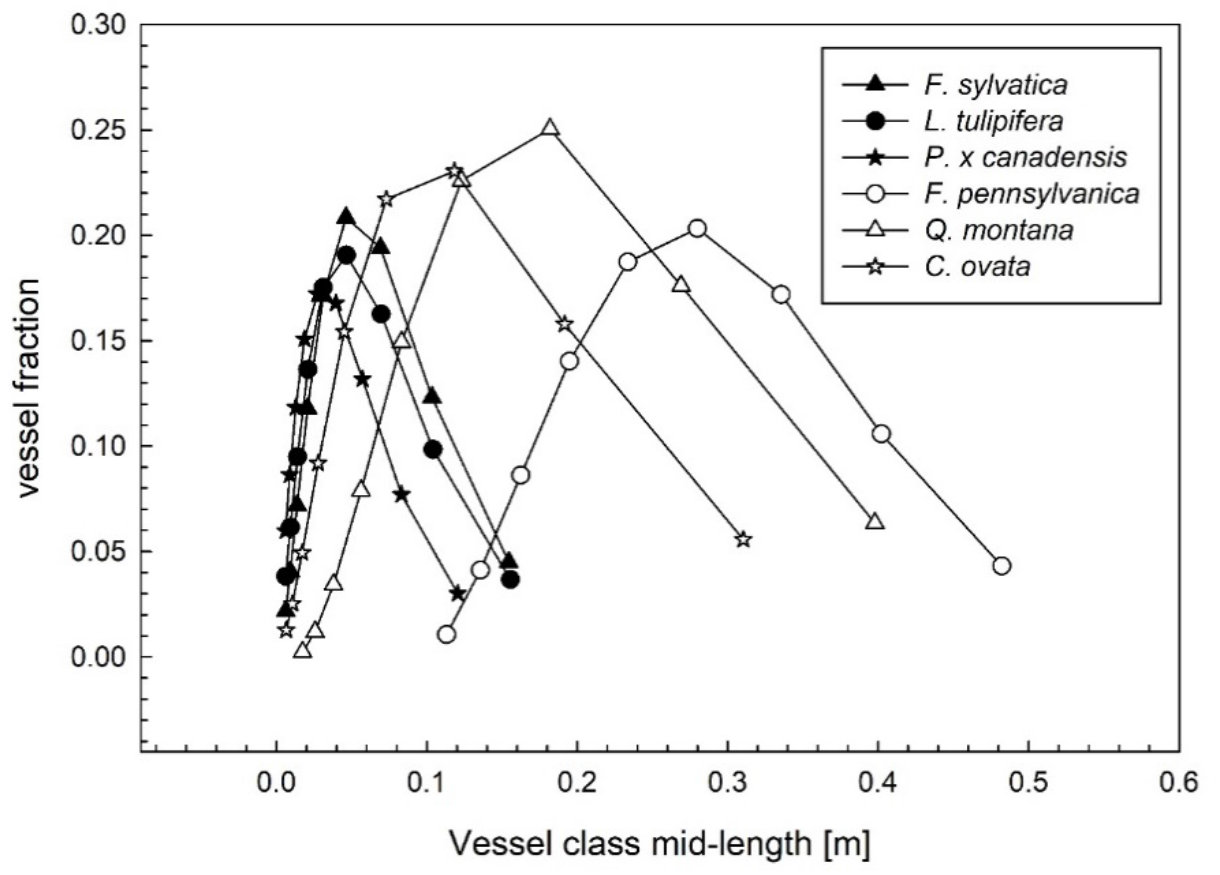
Vessel length distribution of three diffuse-porous (filled symbols) and three ring-porous (unfilled symbols) tree species. Six branches per species were sampled and compiled before fitting vessel distribution according to Christmann *et al*., 2009 ((53)).

**Fig S 2.**
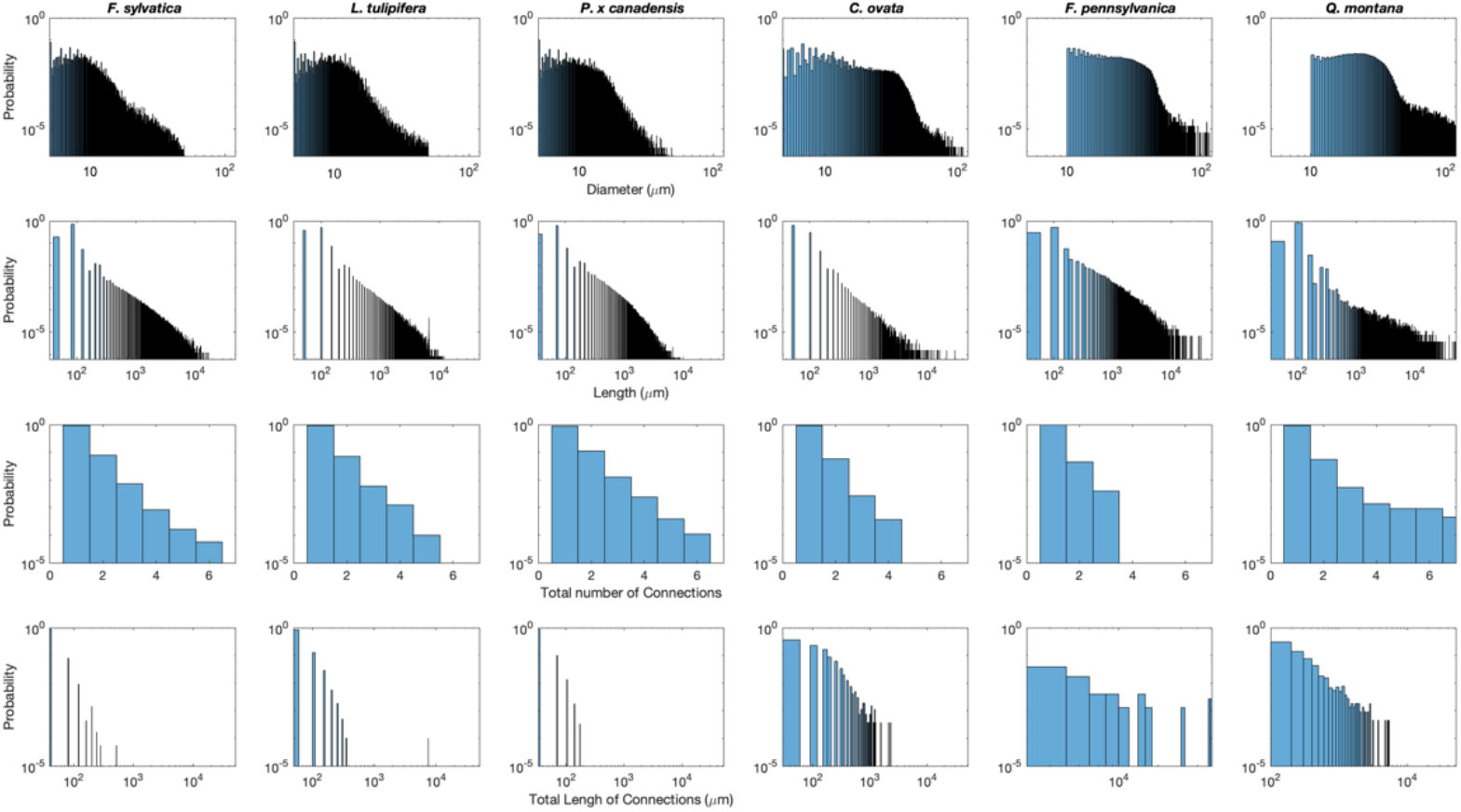
Characteristics of xylem segments and their connections of *Fagus sylvatica, Liriodendron tulipifera, Poulus x canadensis, Carya ovata, Fraxinus pennsylvanica*, and *Quercus montana*. Calculations are based on the ϑ-obrazom network. All data are represented on a log scale with the exception of the total number of connections per xylem segment.

**Fig S 3.**
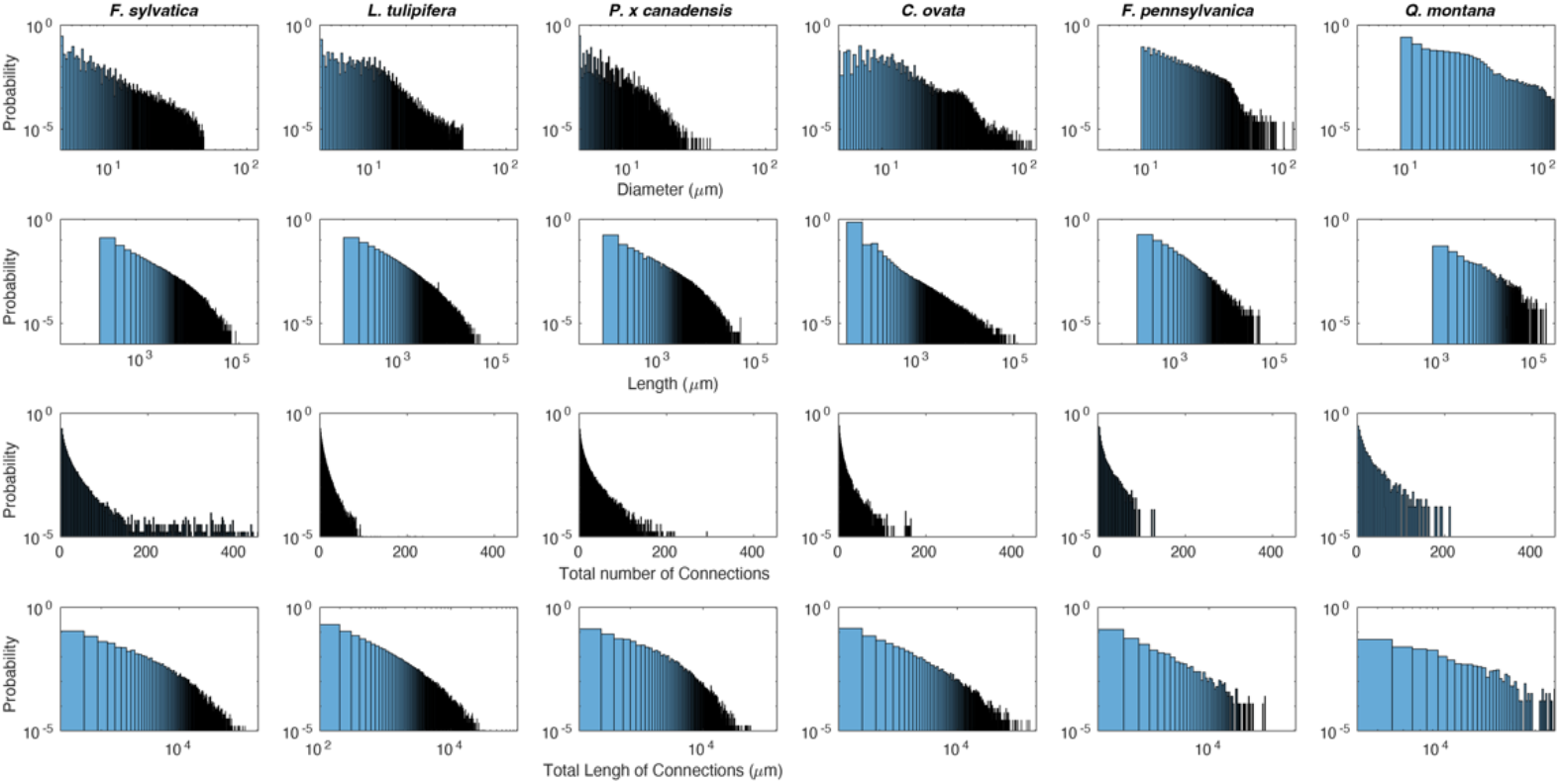
Characteristics of xylem vessels and their connections of *Fagus sylvatica, Liriodendron tulipifera, Poulus x canadensis, Carya ovata, Fraxinus pennsylvanica*, and *Quercus montana*. Calculations are based on the xylem network analysis. All data are represented on a log scale with the exception of the total number of connections per xylem vessel.

**Fig S 4.**
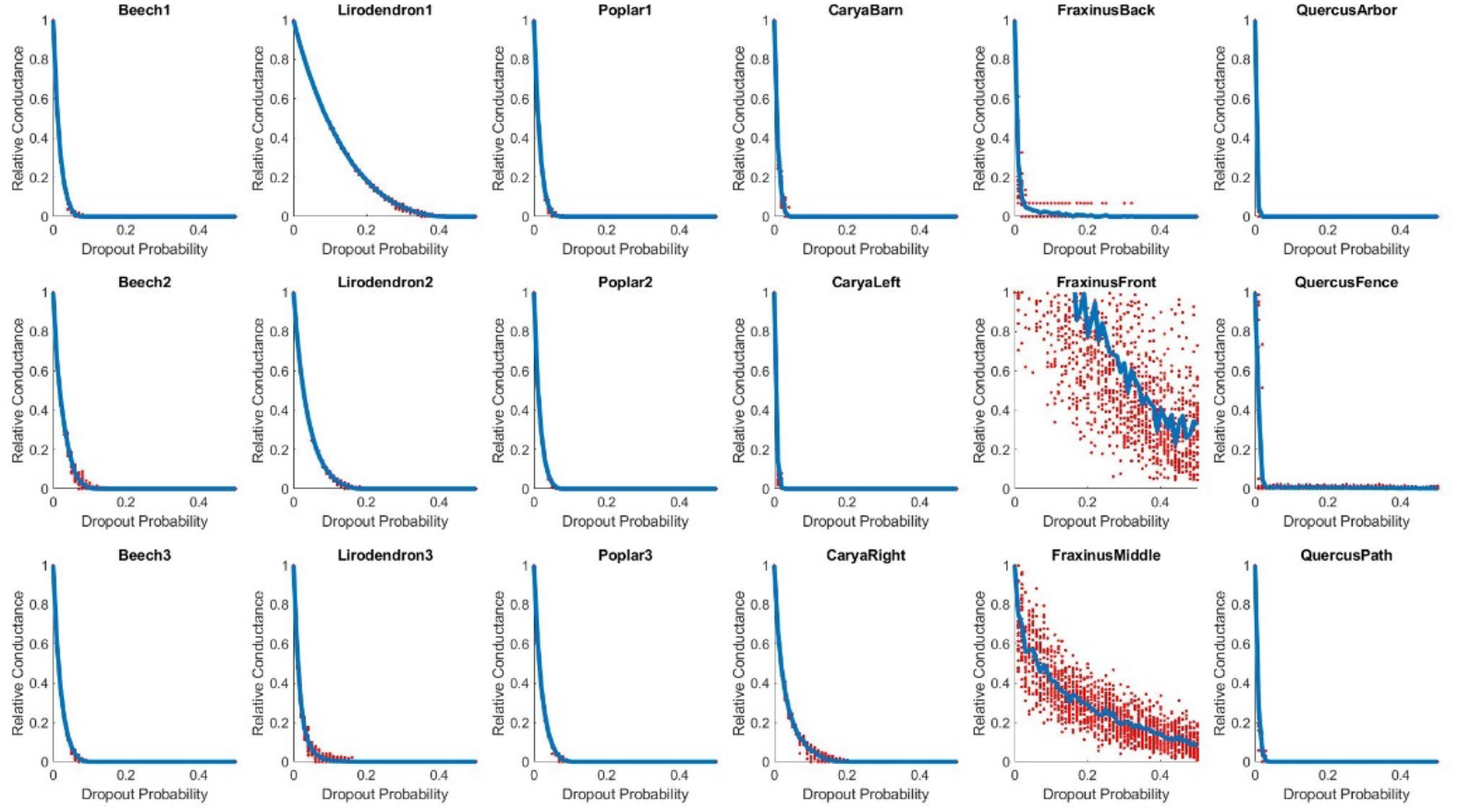
Relation between relative conductance of xylem networks of *Fagus sylvatica* (A), *Liriodendron tulipifera* (B), *Poulus x canadensis* (C), *Carya ovata* (D), *Fraxinus pennsylvanica* €, and *Quercus montana (F)* and increasing vessel dropout probability in fluid dynamic simulation. Each species is represented by three individual samples (three panels). Blue line represents the fit to thirty rounds of simulation in which vessels were randomly excluded.

**Fig S 5.**
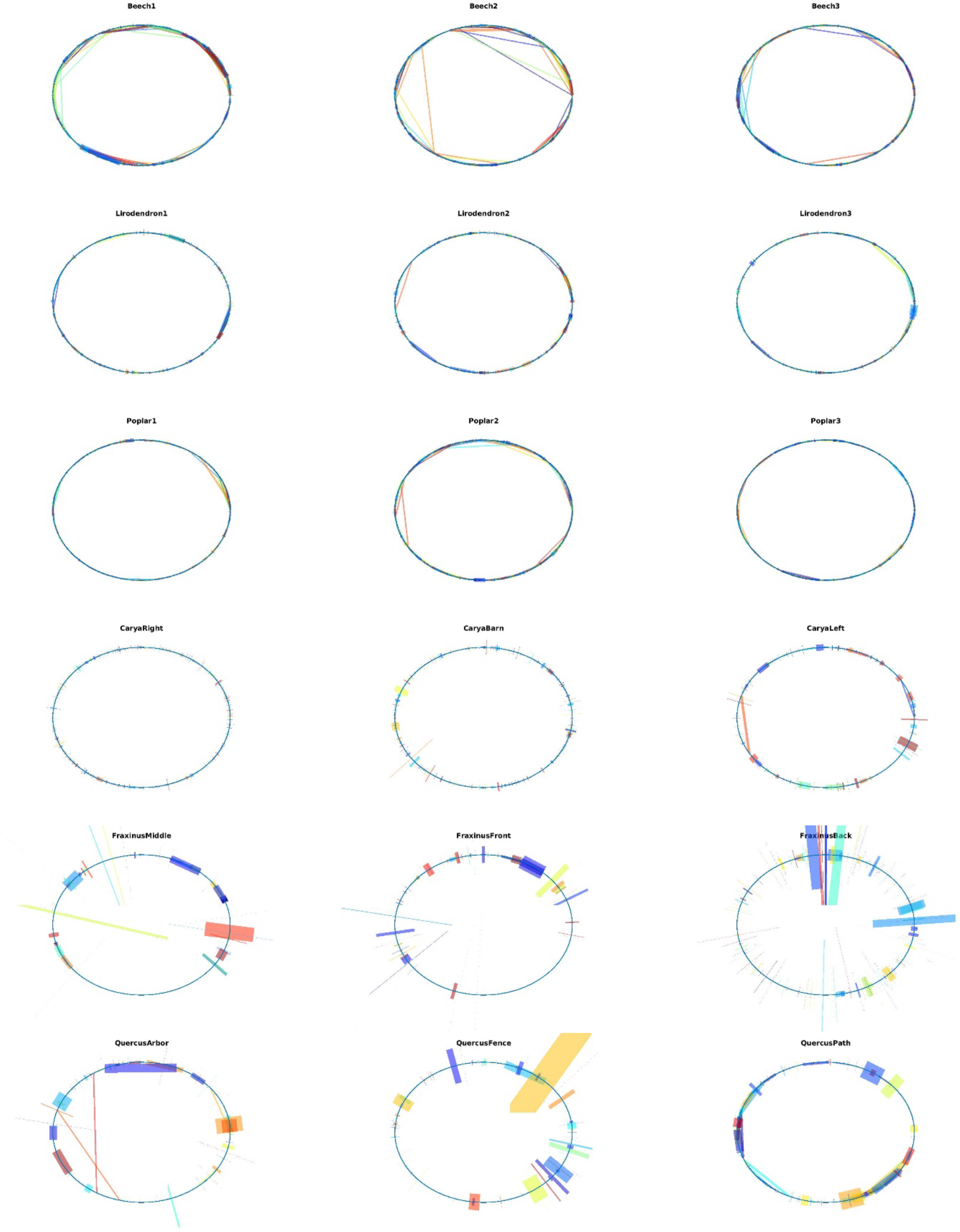
Network presentations of *Fagus sylvatica* (a), *Liriodendron tulipifera* (b), *Poulus x canadensis* (c), *Carya ovata* (d), *Fraxinus pennsylvanica* (e), and *Quercus montana (f)* based on ϑ-obrazom network analysis. Circle graphs can be read as described in Methods and Results. Each species is represented by three individual samples (three panels).

**Fig S 6.**
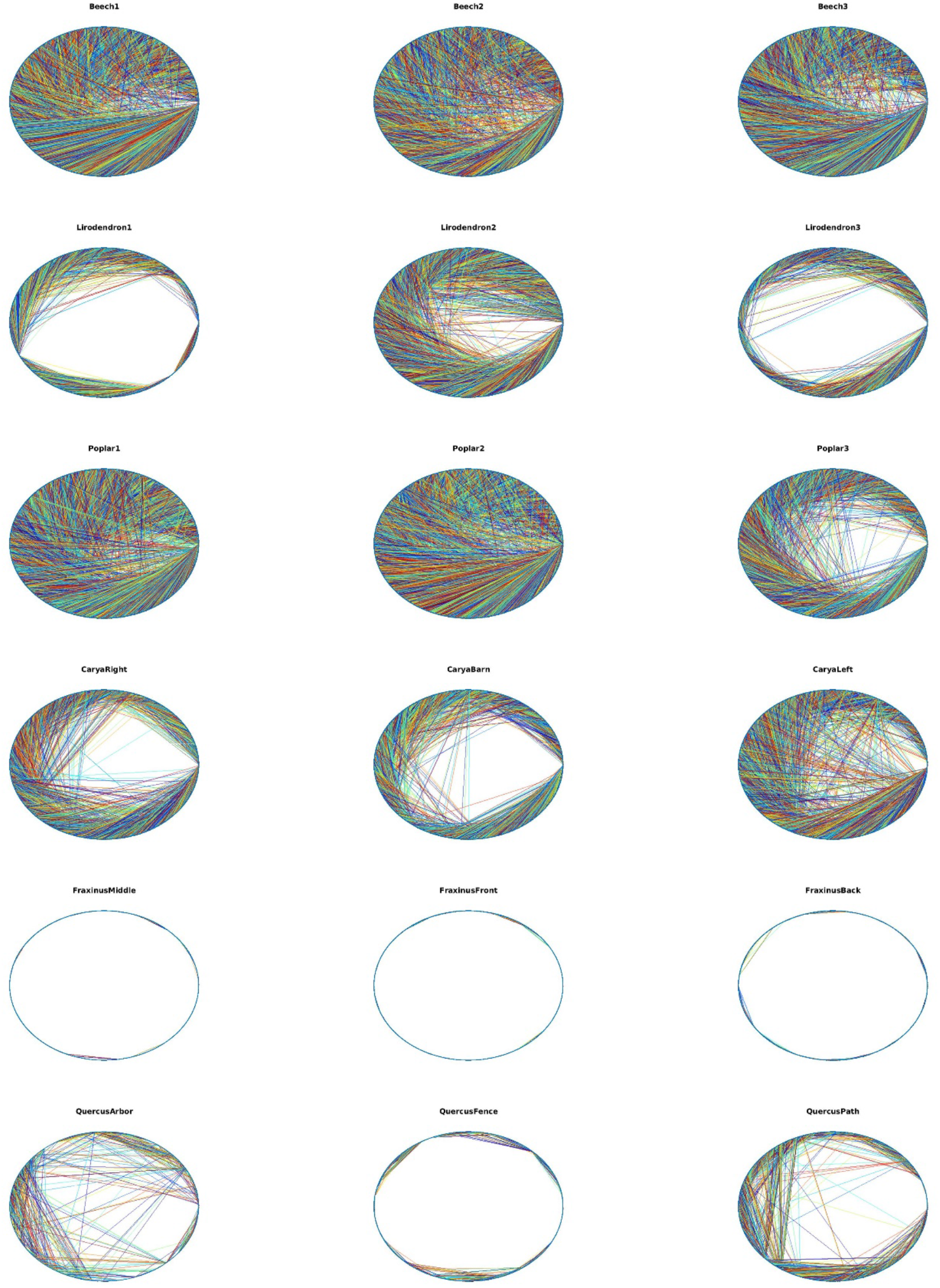
Network presentations of *Fagus sylvatica* (a), *Liriodendron tulipifera* (b), *Poulus x canadensis* (c), *Carya ovata* (d), *Fraxinus pennsylvanica* (e), and *Quercus montana (f)* based on xylem network analysis. Circle graphs can be read as described in Methods and Results. Each species is represented by three individual samples (three panels).

